# Brain network dynamics fingerprints are resilient to data heterogeneity

**DOI:** 10.1101/2020.01.26.920637

**Authors:** Tommaso Menara, Giuseppe Lisi, Fabio Pasqualetti, Aurelio Cortese

## Abstract

Large multi-site neuroimaging datasets have significantly advanced our quest to understand brain-behaviour relationships and to develop biomarkers of psychiatric and neurodegenerative disorders. Yet, such data collections come at a cost, as the inevitable differences across samples may lead to biased or erroneous conclusions. Previous work has investigated this critical issue in resting-state functional magnetic resonance imaging (rs-fMRI) data in terms of effects on static measures, such as functional connectivity and brain parcellations. Here, we depart from prior approaches and utilize dynamical models to examine how diverse scanning factors in multi-site fMRI recordings affect our ability to infer the brain’s spatiotemporal wandering between large-scale networks of activity. Building upon this premise, we first confirm the emergence of robust subject-specific dynamical patterns of brain activity. Next, we exploit these individual fingerprints to show that scanning sessions belonging to different sites and days tend to induce high variability, while other factors, such as the scanner manufacturer or the number of coils, affect the same metrics to a lesser extent. These results concurrently indicate that we can recover the unique trajectories of brain activity changes in each individual, but also that our ability to infer such patterns is affected by how, where and when we try to do so.

**Author summary:** We investigate the important issue of data heterogeneity in large multi-site data collections of brain activity recordings. At a time in which appraising the source of variability in large datasets is gaining increasing attention, this study provides a novel point of view based on data-driven dynamical models. By employing subject-specific signatures of brain network dynamics, we find that certain scanning factors significantly affect the quality of resting-state fMRI data. More specifically, we first validate the existence of subject-specific brain dynamics fingerprints. As a proof of concept, we show that dynamical states can be estimated reliably, even across different datasets. Finally, we assess which scanning factors, and to what extent, influence the variability of such fingerprints.

## Introduction

Untangling the brain’s dynamics at rest is a central aspect in the quest to reveal the mechanisms underlying the spontaneous wandering of the mind between well-established, large-scale networks of neural activity [1–3]. The characterization of the brain dynamics’ spatiotemporal organization into networks has greatly benefited from the creation of very large neuroimaging datasets [4, 5], such as the Human Connectome Project (HCP) [6,7], the UK Biobank [8], and, in the context of neurodegenerative diseases, the Alzheimer’s Disease Neuroimaging Initiative [9]. Large neuroimaging datasets have, furthermore, played a crucial role in the development of novel biomarkers for psychiatric and neurodegenerative disorders [10–12]. Yet, appraising how differences in physical parameters or scanning protocols affect the quality of these data – especially fMRI recordings – remains an outstanding problem [13–16]. For instance, imaging sequences are considerably affected by site-dependent differences such as scanner drift over time, or maintenance routine [16]. Only few recent works have addressed the problem of data variability in rs-fMRI data across sites [17–20], while some others have proposed techniques to harmonize multi-site data [10, 12, 14–16, 21, 22]. Despite growing interest in the intricacies inherent to multi-site data, this line of research is still in its infancy (the first publication appeared in 2013 [23]). Furthermore, although the brain is a complex dynamical system capable of exhibiting rich nonlinear dynamics [24, 25], most studies to date have relied on static measures (e.g., functional connectivity); and little to no attempts exist at exploring such issues from the viewpoint of dynamical models.

Data-driven dynamical models are a promising and powerful tool for the analysis and prediction of the spatiotemporal organization of brain activity [26–29]. These models allow us to harness the vast amount of spurious information contained in large datasets [30–32], capture the hierarchical organization of brain activity [33], enhance brain-computer interfaces [34, 35], and may even be employed in clinical settings [10, 36–38]. However, how the inference and identification of dynamical models is affected by different factors in multi-site data acquisition has yet to be investigated. Additionally, dynamical models could provide fine-grained insight into the extent of the effect of these factors on the data.

One limitation of data-driven models is that, generally, large amounts of data are needed to train the model in the first place. Here, we avoid this issue by employing two datasets. We leverage the high number of subjects (*n*_HCP_ > 1000) with rs-fMRI data available in the HCP dataset [6], to train a stable and reliable Hidden Markov Model (HMM). An HMM infers brain network dynamics from rs-fMRI time series, where networks are probability distributions representing graphs. We then apply the pretrained HMM to the smaller (*n*_TS_ = 9) Traveling-subject dataset, which consists of a novel, state-of-the-art collection of rs-fMRI measurements of nine healthy subjects who traveled to twelve different sites and were scanned under various conditions (different sites, days, phase encoding, number of channels/coils, manufacturer, scanner; see Materials and Methods and Table S1 for a full list of scanning factors and attributes) [22]. This way, we were able to infer subject-specific brain states and investigate how the retrieval of brain state time courses is affected by an array of scanning factors. Training the model on the HCP data guaranteed that (i) the model was inferred on a large sample, made of carefully collected and homogeneous data and that (ii) the model was stable and did not over-fit on a dataset of limited size. We illustrate the methodological approach in Fig. 1.

**Fig 1.**
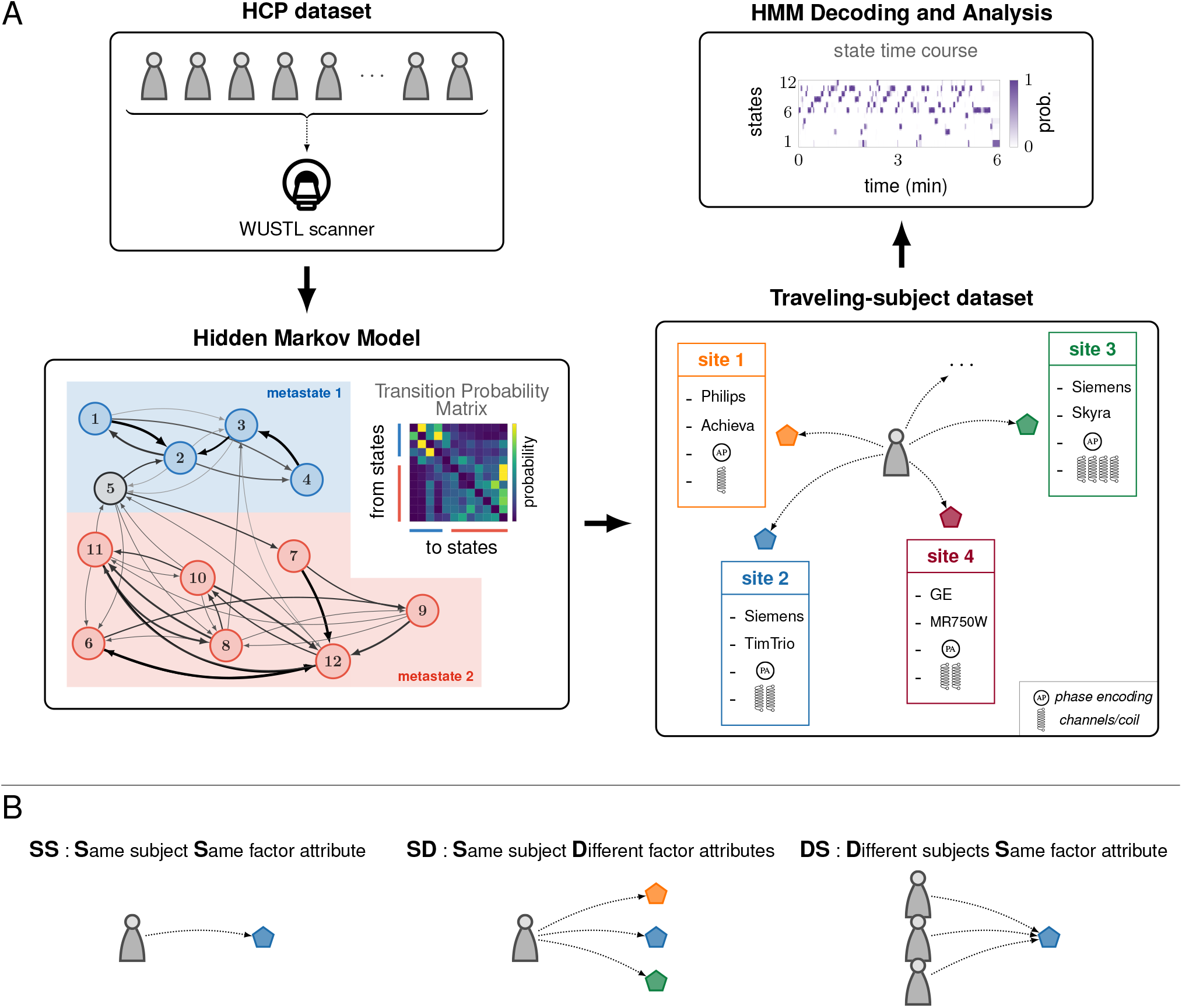
Conceptual flow of the analysis and modeling approach. **A**, rs-fMRI data from the HCP dataset, collected at the Washington University in St. Louis (WUSTL) Connectome-Skyra scanner, were used to infer a Hidden Markov Model (HMM). This model is described by a transition probability matrix, which encodes the probabilities of jumping from one state to another at each time step. Following [33], 12 states were identified and the graph depicted in the figure illustrates the largest transition probabilities (> 0.1) in our model (see also Fig. S2). The states are color-coded in order to distinguish which set of strongly connected states (*metastate*) they belong to. HMM decoding was then applied to the Traveling-subject dataset, in which rs-fMRI data was collected from subjects travelling to different sites. The state time courses from the Traveling-subject dataset were finally used to (1) validate the subject-specific fingerprints associated with states’ dwelling probabilities and the 2-metastate structure put forth previously [33], and (2) analyze the impact of different factors, e.g., site, or scanner model, on fMRI measurements. **B**, To gauge how different factors influenced fMRI data collection, the state time courses obtained from the HMM decoding procedure were compared within and across three different groups: SS (Same subject Same factor attribute), SD (Same subject Different factor attributes), and DS (Different subjects Same factor attribute). In this panel, these three categories are illustrated for the factor *site*, whose attributes consist of the different geographical locations.

Thus, we first utilize the trained HMM to validate the findings on rs-fMRI fingerprints – robust and reproducible quantitative signatures – reported in previous work [28, 33, 39]. We then generalize these findings by applying the HCP-trained HMM to Traveling-subject dataset. This important step allows us to exploit the HMM to assess if, and to what extent, mixed scanning factors affect subject-specific fingerprints and, thus, rs-fMRI recordings. We depart from previous work, which has mostly relied on static functional connectivity / correlation measures and smaller datasets, by exploiting dynamical brain network collective states at a finer temporal resolution. Altogether, this paper juxtapose complementary, yet contrasting, results with respect to rs-fMRI data analysis: we confirm previous findings reporting subject-specific fingerprints, but we also shed light on the presence of factors that induce variability in such fingerprints and, thus, the homogeneity of multi-site fMRI data collections and subsequent inference from the viewpoint of dynamical models.

## Results

In this work, we utilized Hidden Markov model(s) (HMM) to capture the dynamical evolution of brain states in subjects scanned at rest. In neuroscience and neuroimaging, HMMs are typically used to represent the stochastic relationship between a finite number of hidden states that underlie the brain’s complex dynamics, whose evolution in time is captured by the measured data.

First of all, we inferred a single HMM from the high number of individual rs-fMRI recordings collected at a single site that are available in the HCP dataset (*n*_HCP_ = 1003 subjects). To homogenize the input data, the HMM inference was performed on normalized (zero mean and unitary standard deviation) 50-dimensional time series obtained by spatial group Independent Component Analysis (ICA). All subjects’ independent components time series were concatenated along the time dimension, so that the HMM was estimated on this data with 12 states (the number of states was chosen based on previous work [33]). Next, the smaller Traveling-subject data was processed in order to match the HCP time series spatially (by using the HCP ICA spatial maps to project the Traveling-subject data to the same 50 components space of the HCP time series) and temporally (by up-sampling the data to match the sampling rate of the HCP time series, so as to not violate the temporal definition of the model). See Materials and Methods for a detailed account of data processing and model training. Finally, we applied the HCP-trained HMM model to the processed Traveling-subject time series to answer the following questions: are HMM-estimated brain dynamics robustly subject-specific (fingerprints)? Which scanning factors impact measures of brain dynamics the most? What is the magnitude of their effect?

Briefly, an HMM is described by a Transition Probability Matrix (TPM), which encodes the probability of transitioning from one state to another at each time step. The temporal characteristics of such states are expressed as the Fractional Occupancy (FO) – the fraction of time the brain spent in each state. FOs can be computed for each state, subject, or scanning session. Stacking the Fractional Occupancy values of all subjects for the same brain state yielded a vector of FOs whose length corresponded to the number of subjects in the dataset. The pairwise correlation of these vectors generated a *(no. of states)* × *(no. of states)* matrix, named FO Correlation Matrix, defined as

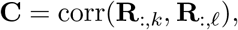

where **R**_:,*k*_ denoted the column vector of the FO of all subjects for the *k*-th state. This matrix captured the overall organization of brain dynamics across states, and its entries quantified the affinity between the FOs of each pair of states across all subjects. In other words, the FO Correlation Matrix highlighted the similarities and dissimilarities between brain states, and encoded the temporal characteristics of brain network dynamics. The organization of the FO Correlation matrix’ revealed (both by visual inspection and by numerical investigation) the emergence of two groups of states – metastates – in which brain state time courses tended to remain. That is, metastates are distinct sets of functional network states that the brain has a propensity to cycle within.

We made use of the information encoded in the FO Correlation Matrix to calculate two different subject-specific metrics in the Traveling-subject data that were key in this study: the Metastate Profile (MP) Differences and the Fractional Occupancy (FO) Correlations. Loosely speaking, the former provided the difference between the time spent in the two distinct metastates that emerged in our model, compatibly with previous findings [33]. The latter provided the pairwise correlation between the FO vectors of different scanning runs. Due to the availability of numerous scanning sessions for each subject, both metrics could be computed not only across different subjects, but also at the individual level. We capitalized on the robustness of the HMM model inferred on HCP homogeneous data to reveal, through MP Differences and FO Correlations, temporal information of brain state time series in the heterogeneous Traveling-subject dataset. These metrics allowed us to perform a richer analysis than simply limiting ourselves to the study of a model’s TPM – in this context it was one single matrix valid for all subjects.

### Test-retest reliability of brain dynamics estimation

We first inferred the HMM by leveraging the large amount of rs-fMRI data in the HCP dataset. Due to the stochastic nature of the HMM inference – which is based on the probabilistic process of Bayesian inference – the results might vary at each new model training. Thus, we inferred multiple models and selected for further analyses the one with the best fit. Specifically, we inferred *N* = 50 models from random initializations, multiple priors, and different combinations of the available datasets (HCP concatenated with Traveling-Subject, HCP only, and Traveling-Subject only). Next, we ranked the models based on their free energy (an approximation of how well a model fits the data, Materials and Methods). This approach further confirmed the stability of the inference process (see Fig. S1). Because the MP Difference was one of the individual fingerprint/signature measures considered in this study, we also verified that the selected model presented the clearest metastate structure. For each model, we therefore computed the Euclidean distance from the ideal FO Correlation Matrix (Fig. S2) – a matrix that described perfectly correlated FOs within the same metastate. The Euclidean distance from the ideal FO Correlation Matrix gauged how well the metastates emerged in the model’s FO Correlation Matrix. We show in Fig. 2 the HMM selected and employed in this work, which is the model that ranked best with respect to free energy, displayed the smallest distance from the ideal FO Correlation Matrix, and was trained solely on HCP time series.

**Fig 2.**
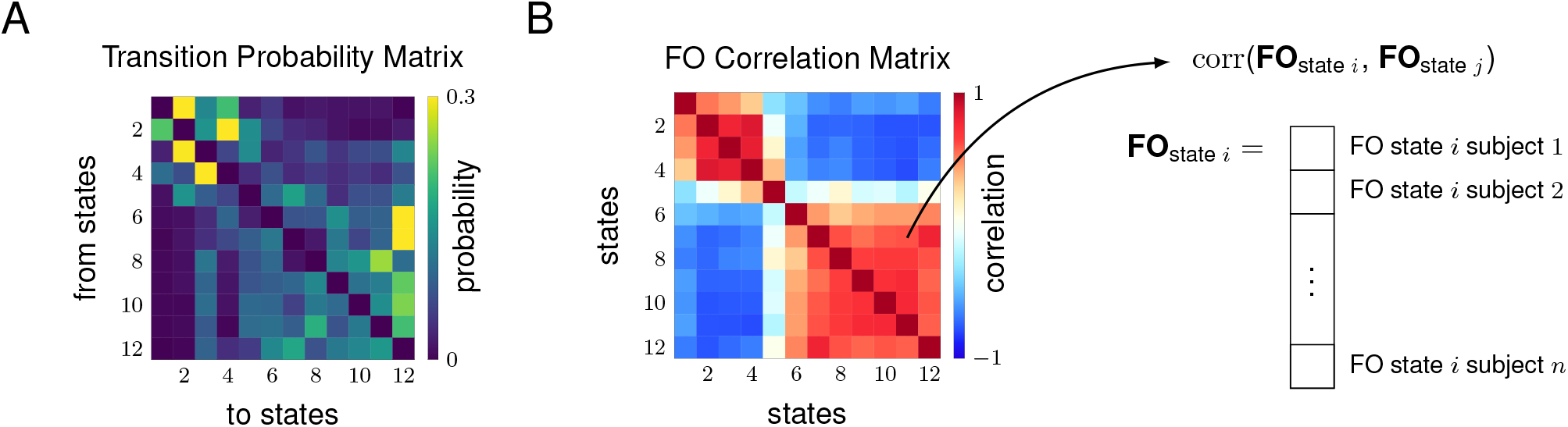
HCP-trained Hidden Markov Model. **A**, Transition Probability Matrix. The emergence of the two metastates can be recognized by simple visual inspection, and was confirmed by a community-detection algorithm. **B**, FO Correlation Matrix. The two metastates are clearly delineated, with state 5 being mostly uncorrelated from all other states [33]. The state FOs are highly correlated (Person correlation > 0.8) within the two metastates across subjects.

Given the stochastic nature of the Variational Bayes approach used to infer the HMM [31], it was unlikely that one would obtain an exact replica of the model originally reported in [33]. However, as displayed in Fig. 2*B* and Fig. S1, all models showed a clear 2-metastate structure, validating the claims that resting-state brain dynamics tend to be hierarchically organized in two larger sets of states (one associated with higher-order cognition, and the other one with sensorimotor and perceptual states, as originally reported in [33]). Moreover, a visual inspection of the TPM matrix alone suggested the emergence of two groups of states that tended to be more (statistically) connected. We confirmed this hypothesis by employing the generalized Louvain algorithm [40] for the discovery of communities in networks.

To note, as mentioned above, we also used the HCP-derived TPM as a prior to train an HMM on the Traveling-subject dataset alone. This choice of prior ensured that the inference started from established initial conditions before dealing with the small size of the Traveling-subject dataset. Surprisingly, although the number of subjects in the Traveling-subject dataset was much smaller than the number of subjects in the HCP dataset, the 2-metastate structure still emerged in the model’s matrices (Fig. S3*D*), as also confirmed by the generalized Louvain algorithm. This result highlighted that, notwithstanding mixed scanning protocols and small sample, the metastates could be retrieved and unfold as a robust feature of resting-state data.

### Metastate Profiles and Fractional Occupancies Are Robust Subject-Specific Fingerprints

Previous findings reported that brain dynamics is subject-specific and nonrandom. To extend this notion, we applied the best-fitting, HCP-trained HMM, to the Traveling-subject dataset, obtaining the state time courses for each 10-minute scanning session. Next, from each individual’s state time courses, we calculated the FO of all 12 states, as well as the Metastate Profile (MP). For a subject *i* and scanning run *k*, MP_*i,k*_ was computed as the FO of the second metastate (states 6-12) minus the FO of the first metastate (states 1-4). We excluded state 5 from our analysis as it was uncorrelated from the other states, had the highest variance, and was previously shown to be associated with head motion in the scanner [33]. As a means to compare different scanning sessions within subjects, or for later analyses, different different factors or subjects, we computed MP Differences and FO Correlations across runs. Specifically, the former was defined as the absolute difference between the Metastate Profiles of different runs; larger values indicate higher variability. For instance, the MP Difference between run *k*_1_ for subject *i*_1_ and run *k*_2_ for subject *i*_2_ read as

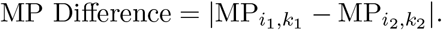

Instead, the FO Correlation between run *k*_1_ for subject *i*_1_ and run *k*_2_ for subject *i*_2_ was defined as

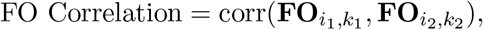

where **FO**_*i,k*_ denoted the 11-dimensional column vector of the FOs of all 12 states minus state 5 for subject *i* and run *k*. We summarize the derivation of these two measures in Fig. 3*A*. To note, here we use the notation subject *i*_1_ and *i*_2_ for a general case, but this naturally applies to two different scans belonging to the same subject (i.e., within-subject comparison).

**Fig 3.**
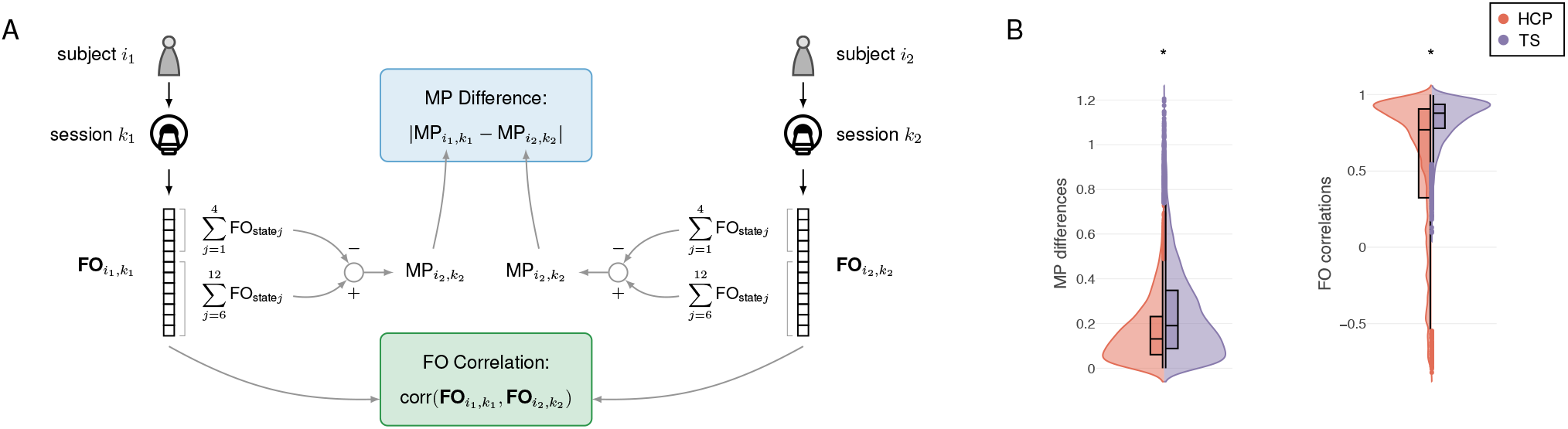
Metastate Profile Differences and FO Correlations computation, and within-subject comparison. **A**, Schematic illustrating the computation of the MP Differences and FO Correlations. To note, subject *i*_1_ and *i*_2_ can mean both the same subject’s data but from different scans, or different subjects. **B**, Comparison of within-subject MP Differences and FO Correlations between the HCP (in red) and the Traveling-subject (TS, in purple) datasets. For MP Differences (left panel) in the HCP data: median = 0.131, lower quartile = 0.61, upper quartile = 0.231; for MP Differences in the TS data: median = 0.191, lower quartile = 0.088, upper quartile = 0.348. For the FO Correlations (right panel) in the HCP data: median = 0.77, lower quartile = 0.324, upper quartile = 0.906; for FO Correlations in the TS data median = 0.876, lower quartile = 0.779, upper quartile = 0.936.

Before delving into the main analyses of the Traveling-subject dataset, we considered the consistency of these two measures of brain activity dynamics both in the HCP and the Traveling-subjects datasets. In both datasets, there were multiple scans per subject (*m*_HCP_ = 4 and *m*_TS_ > 42, respectively), allowing us to compute MP Differences and FO Correlations within subjects. Given the high homogeneity of the HCP dataset, we expected this to provide a lower bound in terms of dissimilarity between scans belonging to a given subject. Strikingly, both datasets had highly similar distributions for MP Differences and FO Correlations (2-sample Kolmogorov-Smirnov test, *k* = 0.18 and *p* < 10^−5^ for MP Differences, *k* = 0.29 and *p* < 10^−5^ for FO Correlations), with low differences (peak = 0.04 for HCP data and peak = 0.06 for TS data, Fig. 3*B* left plot) and high correlations (peak = 0.88 for HCP data and peak = 0.93 for TS data, Fig. 3*B* right plot). These results provided initial (strong) evidence for the presence of – and our ability to infer – subject-specific brain dynamics patterns.

We next interrogated in detail the Traveling-subject dataset. Each scanning factor considered in this study had multiple distinct attributes. For instance, for the factor *scanner manufacturer* there were sessions recorded through scanners produced by three different manufacturers (Siemens, Philips, and General Electric, see also Table S1). We computed the values of MP Differences and FO Correlations for all the runs of the same subject and the same factor attribute (SS), the same subject and different factor attributes (SD), and different subjects but the same factor attribute (DS). A 1-way ANOVA on the median MP Differences (Fig. 4*A*), and on the median FO Correlations (Fig. 4*B*), resulted in a highly significant main effect of comparison group (SS, SD, DS), for both measures (MP differences: *F*_2,15_ = 7.64, *p* = 0.005; FO correlations: *F*_2,15_ = 19.76, *p* < 10^−3^). Applying post-hoc comparisons, we found that, on average, the median MP Differences for the same subject within the same factor (SS) were significantly lower than the median MP Differences for different subjects within the same factor (DS), (~ 38% lower, 2-sided *t*-test, *t*_10_ = −3.59, *p* = 0.005, Fig. 4*A*). Analogously, on average, the median FO Correlations within the same factor for the same subject were higher than across different subjects (~ 10% higher, 2-sided *t*-test, *t*_10_ = 8.15, *p* < 10^−3^, Fig. 4*B*). Strong evidence for how the state time courses of a given subject (within the same factor attributes) tended to be particularly similar was also evident in the FO Correlations median values being greater than 0.9 (group SS, Figs. 3*B* and 4*B*). These findings support the hypothesis that MP Differences and FO Correlations are robust subject-specific measures, as they were resilient to the single effect of all the factors considered in this study.

**Fig 4.**
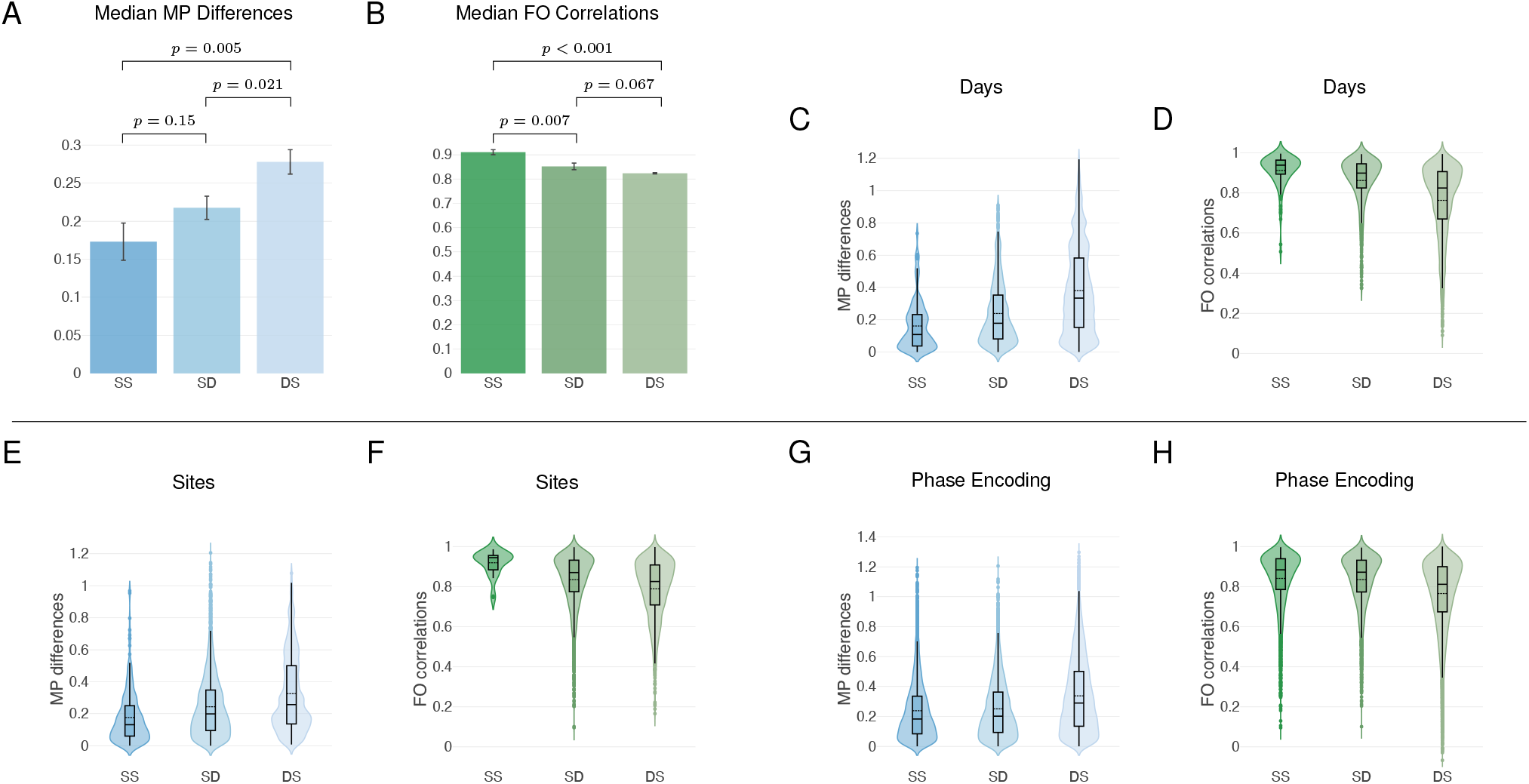
Metastate Profile Differences and FO Correlations within vs between subjects, across scanning factors. **A-B**, The average median of the MP Differences and FO Correlations for the three sets SS (Same subject Same factor attribute), SD (Same subject Different factor attributes), and DS (Different subjects Same factor attribute). MP Differences are the absolute difference between the Metastate Profiles of different runs, while FO Correlations are the pairwise correlation between the Fractional Occupancy vectors of different runs. The set SS consistently displays lower MP Differences and higher FO Correlations than the set DS, confirming the fact that such metrics are subject-specific. The fact that the set SD lies between SS and DS suggests that some scanning parameters influence the aforementioned metrics for resting-state scans of the same subject, but not as much as inter-individual differences. Bars represent the median, error bars the SEM. Statistical comparisons were performed with 2-sided t-tests. **C-H**, Distributions of values for both metrics and all subjects pooled. The set SS comprises the MP Differences (resp., FO Correlations) computed for each subject within the same factor attribute (e.g., for ‘Days’, day 1), and the SS distribution displays these values for all subjects; the set SD consists of the MP Differences (resp., FO Correlations) computed for each subject across different attributes of the same factor (e.g., all possible combinations within ‘Days’), and the SD distribution displays these values for all subjects; finally, the set DS consists of the MP Differences (resp., FO Correlations) computed across all subjects within the same factor attribute, and the DS distribution displays these values for all attributes of the same factor. For all the distributions, the black dashed lines illustrate the mean. In panels **G** and **H** the difference between SS and SD distributions was not significant (Table).

To further substantiate these results, we used a simple machine learning approach to predict individuals based on their brain dynamics fingerprints. We applied logistic regression to classify the individuals in the Traveling-subject dataset by a leave-one-attribute-out cross-validation procedure (Materials and Methods). In brief, for each factor, we repeated the training and validation of the classifier as many times as the number of factor attributes, using each time the samples of one left-out factor attribute as validation set and the remaining samples from all other attributes as training set. We found the accuracy of the classification to be consistently well above the theoretical chance level (9 subjects: 1/9 ≈ 0.11), scoring on average 0.22 ± 0.02 for the classification based on MPs (a single value for each factor attribute) (*t*-test against chance level, *t*_5_ = 16.4, *p* < 10^−4^), 0.30 ± 0.03 for the classification based on FOs (a length-11 vector for each factor attribute) (*t*-test against chance level, *t*_5_ = 17.62, *p* < 10^−3^), and 0.28 ± 0.02 when using the combined measures (*t*-test against chance level, *t*_5_ = 21.56, *p* < 10^−3^). We report the classification results for each factor in see Table S3 (see also Fig. S8).

### In rs-fMRI Data, Not All Factors are Equal

Given that the Traveling-subject dataset contained a considerable number of factors, we inquired which of these factors, and to what extent, influenced the subject-specific fingerprints defined on the HMM state time courses. Specifically, we asked which factors affected the MP Differences and the FO Correlations most, both within and across subjects. Thus, we compared three different groups (SS, SD, and DS, as illustrated in Fig. 1*B*) of MP Differences and FO Correlations, for six different factors, each containing at least two attributes (see Table S1 for the full list of factors and associated attributes).

Although different runs always carried some variability, some factors heavily influenced the MP Differences and the FO Correlations while others did so only minimally. We summarize the main results of this comparison in Fig. 4 and report the additional ones in Fig. S6. We also report in Table the results of Kolmogorov-Smirnov nonparametric tests between all the distributions of values for the groups of MP Differences and FO Correlations. More in detail, by comparing the distributions of values for both metrics between the sets SS (Same subject and Same factor attribute) and SD (Same subject and Different factor attributes), we found them to be statistically different (*p* < 10^−3^, see Table) for all factors except for the phase encoding direction, as also noticeable in Fig. 4*G*-*H*.

**Table 1.** 2-Sample Kolmogorov-Smirnov test results for MP Differences and FO Correlations. The check-mark indicates that the difference is significant (i.e., the null hypothesis that the samples are drawn from the same underlying continuous population can be rejected at the 5% significance level), and the cross otherwise. All *p*-values have been FDR-adjusted [41] and they all satisfy *p* < 10^−3^ when the null hypothesis is rejected. Test statistics are reported in Table S2. SS: Same subject Same factor attribute. SD: Same subject Different factor attributes. DS: Different subjects Same factor attribute.

| Parameter | MP Differences |  |  | FO Correlations |  |  |
| --- | --- | --- | --- | --- | --- | --- |
|  | SS | SS | SD | SS | SS | SD |
|  | vs | vs | vs | vs | vs | vs |
|  | SD | DS | DS | SD | DS | DS |
| 1. Site | ✓ | ✓ | ✓ | ✓ | ✓ | ✓ |
| 2. Day | ✓ | ✓ | ✓ | ✓ | ✓ | ✓ |
| 3. Phase | ✗ | ✓ | ✓ | ✗ | ✓ | ✓ |
| 4. Channels/Coil | ✓ | ✗ | ✓ | ✓ | ✓ | ✓ |
| 5. Manufacturer | ✓ | ✓ | ✓ | ✓ | ✓ | ✓ |
| 6. Scanner | ✓ | ✓ | ✓ | ✓ | ✓ | ✗ |

The MP Differences of the scanning factors day and site had, on average, not only the smallest median and variance for the values in the set SS, but also the largest median differences with the values in the set DS for the same parameters (Fig. 4*C* and Fig. 4*E*, Kolmogorov-Smirnov test *p* < 10^−3^). Analogously, the FO Correlations of day and site had, on average, not only the largest median and smallest variance for the values in the set SS, but also the largest median differences with the values in the set DS for the same parameters (Fig. 4*D* and Fig. 4*F*, Kolmogorov-Smirnov test *p* < 10^−3^). As we elaborate further below, this is in agreement with previous findings on the variability of functional connectivity across sites [19]. We report in Fig. S5 the distributions of values for the scanning parameters that lied between the aforementioned opposite cases – all significantly different (*p* < 10^−3^). It is worth noting that the median MP Difference for the same subject within the same factor attribute was, according to our data, ~ 26% smaller than between different factor attributes (2-sided *t*-test, *t*_10_ = −1.55, *p* = 0.15); compatibly, the median FO Correlations were, on average, ~ 6.5% higher in the group SS than in the group SD (2-sided *t*-test, *t*_10_ = 3.43, *p* = 0.007).

Finally, the machine learning classification of brain dynamics fingerprints described earlier, were qualitatively generally in agreement with these findings. Leave-one-attribute-out classification revealed that, for both fingerprints, the accuracy in predicting individual subjects was the lowest when the training and validation sets were based on different days (see Table S3).

### Scanning Sites and Days Increase the Variability of Individual Brain State Time Courses

To further evaluate the influence that different scanning variables have on MP Differences and FO Correlations, we directly compared their effects across these fingerprints. We first analyzed the raw medians of the distributions of MP Differences and FO Correlations for each scanning factor in the groups SS (Same subject Same factor attribute), SD (Same subject Different factor attributes), and DS (Different subjects Same factor attribute). We found that, while both fingerprints possessed a shared variance (Fig. 5*A*, Coefficient of determination *R*^2^ = 0.375), they also provided independent information. In fact, MP Differences displayed consistently larger median differences within the three groups of values (SS, SD, DS) than FO Correlations (Fig. 5*A*, 2-sided Wilcoxon signed rank test, *z* = 3.68, *p* < 10^−3^).

**Fig 5.**
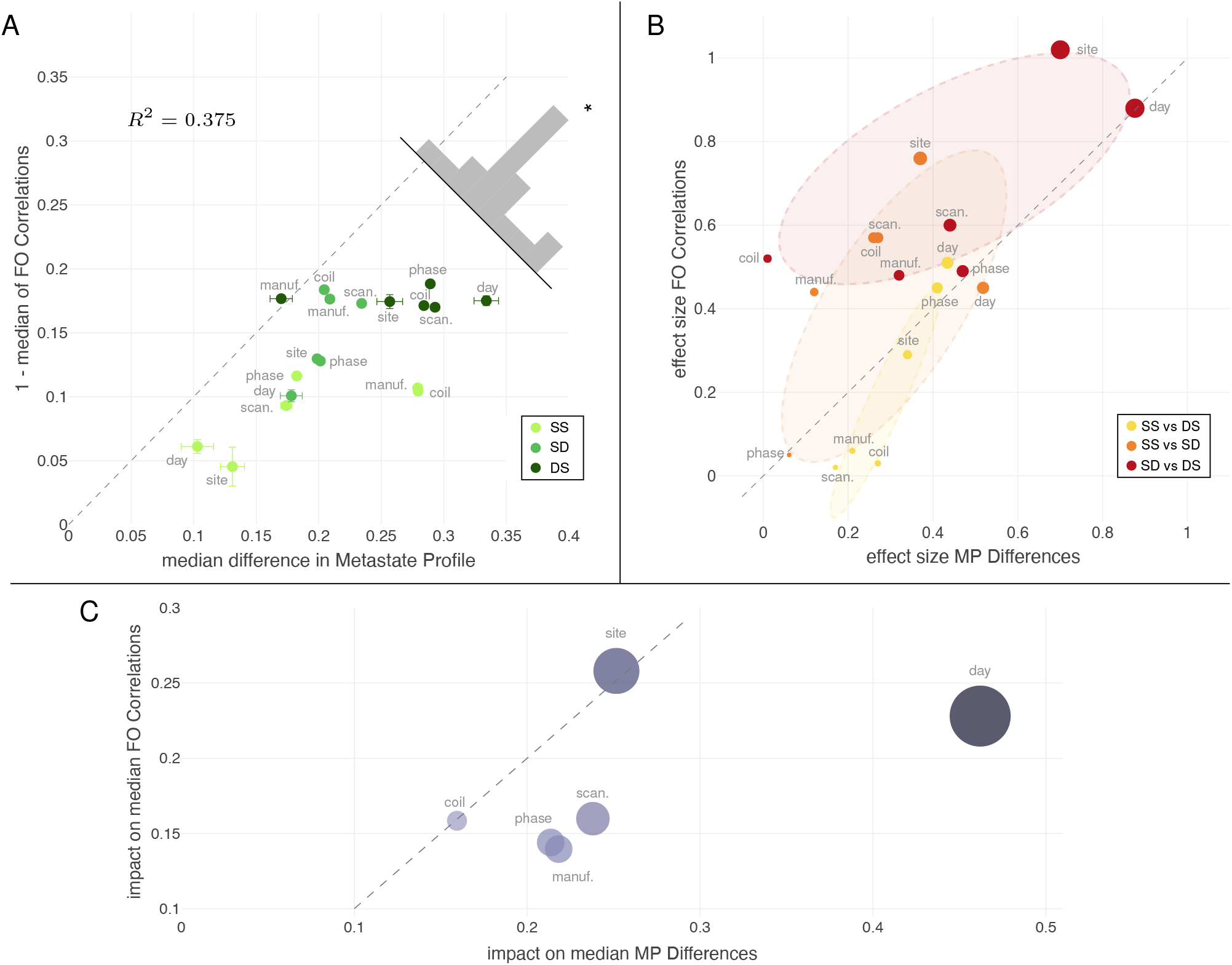
Effect of scanning factors within and across MP Differences and FO Correlations distributions. **A,** The *x* and *y* axes represent the median of MP Differences and 1 - median of FO Correlations, respectively, for different scanning factors in different groups SS, SD, and DS, along with the standard error of mean. The dashed line represents the diagonal *y* = *x*. The median of MP Differences is more affected by all of the scanning factors (Wilcoxon signed-rank test, *z* = 3.68, *p* < 10^−3^). **B**, The effect size (Cohen’s *d*) values obtained by comparing the log-transformed distributions of the MP Differences and FO Correlations across different scanning session factors and attributes. The ellipses represent the least squares minimization of the distance from the cloud of points for each of the three sets [42]. The largest effect on both the metrics is consistently caused by the factors site and day, for all the comparisons between groups of distributions. **C**, The *impact* due to different parameters on the medians of the SS, SD, and DS distributions of values across all different scanning parameters. Again, MP Differences are impacted the most by all scanning factors, especially by site and day.

Next, to get an unbiased estimate of the effect size of each factor on the distributions of MP Differences and FO Correlations, we computed Cohen’s *d* from the log-transformed distributions of the MP Differences and FO Correlations across all scanning factors, between groups SS, SD and DS. Fig. 5*B* highlights how the dissimilarity between brain dynamics fingerprints was the greatest when comparing, for the same scanning factor, measures from the same subject and measures from different subjects. Interestingly, there was a stronger contribution to this dissimilarity from MP Differences when considering within factors cases (SS vs DS), but from FO Correlations when considering across factors cases (SD vs DS and SS vs SD). These results complement, from a dynamical point of view, both seminal and more recent work reporting more dissimilar resting-state networks inter-subject than intra-subject [16, 23, 39, 43].

Finally, we assessed which scanning factors influenced the median values of the three groups SS, SD, and DS the most by defining their *impact*. For a given fingerprint, this impact was calculated as the sum of the absolute value of the differences of medians:

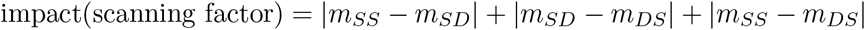

where *m* denoted the median. The scanning factors site and day yielded the largest median differences for both MP Differences and FO Correlations (Fig. 5*C*), with overall effects almost twice as large as those carried by other factors.

In conclusion, site and day were consistently associated with the largest effect sizes in all three (color-coded) groups (Fig. 5*B*). Moreover, site and day were also the co-variate that caused the largest median differences in the distribution values of the three groups (Fig. 5*C*). Conversely, some scanning factors seemed to have a rather small effect on the fingerprints distributions. Such “negligible” scanning factors were the number of channels per coil and the phase encoding direction, which had a limited impact on the metrics considered in this work.

## Discussion

In this work, we addressed the issues of reproducibility and variability of fMRI data from the angle of brain dynamics. We leveraged the large HCP collection of rs-fMRI data to infer a hidden Markov model capable of describing brain state time courses at the subject level. By applying such a model to a dataset of traveling subjects, we found that brain network dynamics displayed signature fingerprints that were robust to different physical and temporal factors affecting the data that populates multi-site collections of neuroimaging data. This study corroborates and complements previous work that found that the emergence of temporal patterns of brain activity tend to repeat more similarly within the same subject and over time [28, 33, 44]. This result promotes further investigations on the dynamical characteristics of brain states.

Recent years have witnessed a growing interest in the identification and characterization of the factors that tend to introduce spurious effects in multi-site fMRI recordings, endangering the reproducibility and the overall quality of the results that may be inferred from these data. The first warnings came from a study that investigated sources of nuisance variation across multiple sites and their impact on rs-fMRI data [23], followed by a number of studies that reported mostly consistent results [14–16,19,20]. Although we used different methods, our finding that different sites are likely to affect fMRI data reliability the most is in line with prior reports [16, 19]. Furthermore, the present study made use of larger and more detailed datasets. The Traveling-subject dataset contains the largest number of subjects out of all the aforementioned studies. To date, only [16] has more sites than the Traveling-subject dataset used in this study, but it has the drawback of scanning only a single subject. Furthermore, the Traveling-subject dataset allowed for the analysis of some scanning factors – such as the numbers of channels per coil or different scanner models within the same vendor – that have not been taken into consideration in previous work, giving more breadth and depth to our findings. Nevertheless, it is important to stress out that the potential variability brought in by scanning factors may not always be a necessarily negative feature. In fact, such variability may even be a powerful test for reproducibility of some findings. For instance, although different scanning factors may be confounds for certain analyses (e.g., comparing participant populations from different sites), they can also be used to test the robustness of a model when generalising analyses across sites with different scanning parameters.

Functional connectivity – typically computed as the correlation between time series representing the average activity in a brain region – has been the mainstay in the analysis of variability in fMRI data. Previous work has demonstrated that a sizeable amount of recordings from the same site enables precise measurements of individual variations in functional connectivity [39], and that individual differences in functional networks are not affected by anatomical misalignment [43]. Here, we complement such studies by showing that individual signatures can still be (easily) recovered within limited recordings from multiple sites (i.e., in the Traveling-subject dataset). To note, the comparison of functional connectivity between scanning sessions is inherently different from the comparison of state time courses. Differently from functional connectivity computed over a whole scanning session, the HMM captures the local temporal wandering of brain activity across states (networks). Therefore, while comparing functional connectivity between different subjects may be akin to comparing longer-term traits, comparing state time courses between subjects may be more closely aligned to comparing a repetition of sequences of brain states at rest.

While the aforementioned studies on functional connectivity have significantly increased our understanding of the brain as a system that obeys network-wide principles, they are mainly agnostic to temporal dynamics within the scanning sessions. This may prevent the level of precision that could at times be the most clinically relevant [45]. Differently from [16, 20], where time seemed to play a negligible effect, we found that different scanning days greatly affected our estimation of brain network dynamics. As mentioned earlier, an intuitive explanation for this apparent discrepancy is that functional connectivity tends to be associated to more coarsely defined subjective traits, whereas an HMM, being inherently more sensitive to temporal differences, is apt to capture more instantaneous cognitive processes. It is worth noting that our findings do not go against the claim that functional connectivity networks remain a reliable subject-specific fingerprint over long period of times, but rather we suggest that brain state trajectories can differ extensively between days, probably due to different cognitive or mental processes. As such, we suggest that dynamic and static measures in fact carry complementary information, which may provide additional insight when used in combination. However, due the limited number of scanning sessions taken on different days in the Traveling-subject dataset (see Table S1), future studies will need to further validate this fact.

How do HMMs compare with the sliding window approach? The sliding window analysis is typically used to improve the temporal definition of functional connectivity studies [46]. Albeit being intrinsically easier to set up, it has crucial limitations. Namely, the sliding window size is constrained by a trade-off between time resolution and quality of the results, and the conclusions from sliding window studies tend to be affected by sampling variability [47]. Conversely, HMM is as fast as the data modality allows, since it provides instantaneous likelihood of high correlation between brain signals [33].

If the goal of a study is a robust and detailed description of a system’s dynamics, the HMM approach requires large amounts of data for training purposes, thus appearing not suitable to analyze small cohorts of subjects. However, in this study we give proof-of-concept that one can use a very large dataset (i.e. HCP) to infer an HMM, which can then be applied to a smaller dataset. Our results indicate that this procedure is robust. Interestingly, if only a relatively small number of subjects is available for the inference process, it is still possible to recover a coarser – and nonrandom – representation of the brain dynamics by using the TPM inferred from a large dataset as a prior (see Fig. S4*D*). Thus, detailed analyses and claims based on hidden Markov modeling should be gauged on the size of the available data. This is a common requirement in neuroimaging studies, as functional connectivity studies also require large amount of data to enable precise measurements [39].

Despite its capabilities, hidden Markov modeling is based on some premises (see also [31, 33] for thorough discussions). It is worth noting that the HMM builds on the Markovian assumption, theorizing that we can predict, based on the state we are at time *t*, which state is more likely to follow at time *t* + 1. Yet, while the brain may violate this assumption due to established long-range temporal dependencies [4], FO Correlations and MP Differences inherently display information that appears at longer time scales.

There are some limitations to this work. For example, while the decoding approach utilized here - training the HMM on the HCP data, and infer brain states trajectories in the Traveling-subject dataset, is a strength of this study, it is also one of its limitations. The HCP and Traveling-subject datasets harbour some differences relating to the scanning protocols or even the countries in which the data were collected (US and Japan). For instance, the sampling rate of the two datasets were originally different (TR_HCP_ = 0.72 s and TR_TS_ = 2.5 s). As such, the up-sampling of the Traveling-subject dataset may have been sub-optimal and thus bias the overall HMM-based brain state dynamics estimation. Yet, the presence of these very differences appear to corroborate the finding that brain dynamics fingerprints are subject-specific. Specifically, we still find that, on average, the MP Differences (resp., FO Correlations) are lower (resp., higher) within subjects than across subjects, even when comparing runs with different scanning parameters. The fact that, at the within-subject level, these two measures had very similar values to those obtained from the HCP dataset (where both model and fingerprints were derived from the same data) provides strong support for this interpretation (i.e., brain dynamics fingerprints are subject-specific), such that it is unlikely that these results are due to inherent bias or noise. A second limitation may arise from the factors that were considered in the Traveling-subject dataset. Although there are several factors, some with many attributes (e.g., there are 12 sites), these factors are sometimes nested within each other. For instance, within the same phase encoding attribute there are scans belonging to different sites. This aspect may have partly influenced (reduced) the effect size of such factors which are heterogeneous with respect to *other* factors, while factors such as day or site would remain unaffected, since these scans were recorded at the same site, with the same protocol.

Given the considerable recent advances in inference techniques [31, 48, 49], and the ever-increasing availability of computational power, our work further suggests that the HMM is, and, most importantly, will be, a powerful technique to explain and interpret the dynamic aspects of the brain. Furthermore, the possibility of inferring an HMM on a very large dataset to apply it to a much smaller one has important implications for clinical applications. In the future, perhaps with even more data, these general models could be built and then utilized to infer subject-specific fingerprints in other smaller cohorts and be used for a more personalized approach to treatments. In other words, a one-size-fits-all approach could be employed to build the model in its general terms, consequently allowing us to move to a personalized course of action by evaluating the model at the individual level. For instance, closed-loop fMRI neurofeedback [50, 51] could significantly benefit from these models, which will allow for a more holistic approach to the dynamical properties of mental and cognitive processes, particularly from a clinical perspective [37, 45, 52, 53].

## Materials and Methods

### Datasets

The two dataset used in this study are (1) the HCP 1200-subject distribution (data available at https://db.humanconnectome.org) and (2) the Traveling-subject dataset (data available at https://bicr-resource.atr.jp/srpbsts/afterfreeregistration). The former consists of rs-fMRI data from *N* = 1206 healthy subjects (age 22-35) that were scanned twice (two 15-minute runs) on two different days, one week apart, on a Siemens 3T Connectome-Skyra scanner. For each subject, in total four 15-minute runs of rs-fMRI time series data with a temporal resolution of 0.72 s and a spatial resolution of 2-mm isotropic were available. For our analysis, we used time series from the 1003 subjects with 4 complete scanning sessions. The HCP dataset provides the required ethics and consent needed for study and dissemination, such that no further institutional review board (IRB) approval is required.

The Traveling-subject dataset consists of 9 healthy subjects (all men; age range 24–32; mean age 27 ± 2.6y), who were all scanned at each of the 12 sites, producing a total of 411 10-minute scanning sessions [22]. Each participant underwent three rs-fMRI sessions of 10 min each at nine sites, two sessions of 10 min each at two sites (HKH and HUH), and five cycles (morning, afternoon, following day, following week, and following month) consisting of three 10-min sessions each at a single site (ATT). In the latter situation, one participant underwent four rather than five sessions at the ATT site because of a poor physical condition. Thus, a total of 411 sessions were conducted [8 participants × (3 × 9 + 2 × 2 + 5 × 3 × 1) + 1 participant × (3 × 9 + 2 × 2 + 4 × 3 × 1)] (see SI Table 1 for all the details on the scanning protocols). In total, there were two phase-encoding directions (posterior to anterior [*P* → A] and anterior to posterior [A → P]), three MRI manufacturers (Siemens, GE, and Philips), four numbers of channels per coil (8, 12, 24, and 32), and seven scanner types (TimTrio, Verio, Skyra, Spectra, MR750W, SignaHDxt, and Achieva). All participants in all datasets provided written informed consent. All recruitment procedures and experimental protocols were approved by the institutional review boards of the principal investigators’ respective institutions (Advanced Telecommunications Research Institute International [ATR] [approval numbers: 13–133, 14–133, 15–133, 16–133, 17–133, and 18–133], Hiroshima University [E-38], Kyoto Prefectural University of Medicine [KPM] [RBMR-C-1098], SWA [B-2014-019 and UMIN000016134], the University of Tokyo [UTO] Faculty of Medicine [3150], Kyoto University [C809 and R0027], and Yamaguchi University [H23-153 and H25-85]) and conducted in accordance with the Declaration of Helsinki.

### Hidden Markov Model

Hidden Markov modeling is a powerful technique that enables the description of time series extracted from a system of interest. Analyses of hidden Markov models seek to recover the sequence of states from some observed data. The underlying assumption of this class of models is that the observed time series of data can be explained by a discrete sequence of hidden states, which must be finite in number. Additionally, to describe a hidden Markov model, an observation model needs to be chosen. We assume multivariate Gaussian observation model, so that, if **x**_*t*_ denotes the data at time step *t*, and *s_t_* represents the state at time step *t*, we can write, whenever state *k* is active,

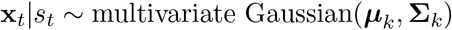

where 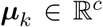 is the vector of the mean blood oxygen level-dependent (BOLD) activation for each channel, with *c* being the number of channels in the data, and 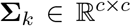 is the covariance matrix encoding the variances and covariances between channels. The transitions between different brain states depend on which state is active at the previous time step. Specifically, the probability of a state being active at time *t* depends on which state is active at time step *t* − 1. This is encoded in the Transition Probability Matrix **Θ**, in which the entry **Θ**_*ij*_ – the transition probability – denotes the probability of state *i* becoming active at the next time step if state *j* is currently active. Formally, by denoting a probability with Pr, it holds that

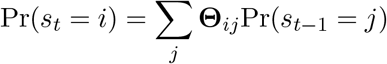

For large datasets, it is possible to resort to stochastic Variational Bayes inference to estimate the posterior distribution of each state (***μ**k*, **Σ**_*k*_), the probability of each state being active at each time step, and the transition probabilities between each pair of states **Θ**_*ij*_ [31]. Finally, notwithstanding the fact that in this study the model has been inferred by concatenating all the subjects – thus implicitly defining the brain states as the outcome of common brain dynamics – the state time courses are subject-specific. That is, the states are inferred at the group level, but the time instants at which each brain state becomes active is subjective and changes between and across subjects.

### Data Preparation and HMM Training

#### HCP dataset

Following [33], extensively preprocessed HCP ICA time series were used for the model training. The preprocessing followed the steps of [6, 54] and is briefly described here. Spatial preprocessing used the procedure described by [55]. Next, structured artifact removal using ICA was followed by FMRIB’s ICA-based X-noisefier (FIX) from the FMRIB Software Library (FSL) [56], which removed more than 99% of the artifactual ICA components in the dataset. Finally, the 50-dimensional extensively preprocessed time series obtained after group spatial ICA are freely available at https://www.humanconnectome.org/study/hcp-young-adult/.

#### Traveling-subject dataset

The dataset was obtained from https://bicr-resource.atr.jp/srpbsts/. Hereafter, we describe the preprocessing procedure that was originally reported in [22]. Raw BOLD signals were preprocessed using SPM8, implemented in MATLAB (R2016b; Mathworks, Natick, MA, USA), The first 10 s of each scan data were discarded to account for T1 calibration. Ensuing preprocessing steps included: slice-timing correction, realignment, coregistration, segmentation of T1-weighted structural images, normalization to Montreal Neurological Institute (MNI) space, and spatial smoothing with an isotropic Gaussian kernel of 6 mm full-width at half-maximum. Thirty-six noise parameters were included in a linear regression model to remove multiple sources of spurious variance (e.g., six motion parameters, average signals over the whole brain, white matter, and cerebrospinal fluid) [57]. Time-series were band-pass filtered using a first-order Butterworth filter [0.01 - 0.08 Hz] to restrict the analysis to low-frequency fluctuations, which are characteristic of rs-fMRI BOLD activity [57]. Finally, to reduce the impact of head motion, scrubbing was performed: framewise displacement (FD) was calculated and volumes with FD > 0.5 mm were removed [58]. Thus, 5.4% ± 10.6% volumes (mean [approximately 13 volumes] ± 1 SD) were removed per 10 min of rs-fMRI scanning (240 volumes). If the number of volumes removed after scrubbing exceeded the average of –3 SD across participants, the sessions were excluded from the analysis. As a result, 14 sessions were removed from the dataset.

Before combining the HCP time series and the Traveling-subject time series for the model inference, we matched the temporal resolution of the two datasets. Specifically, for all results reported in the main text, the Traveling-subject time series were up-sampled in order to match the same repetition time as the HCP data (from TR = 2.5 s to TR = 0.72 s). We also down-sampled the HCP data from TR = 0.72 s to TR = 2.5 s to match the Traveling-subject repetition time. However, the model inferred on HCP down-sampled ICA time-series was not satisfactory, as it did not display a clear 2-metastate structure (Fig. S5). Therefore, we have chosen to re-sample the Traveling-subject data instead of down-sampling the HCP data.

The HMM inference was performed on 50-dimensional standardized ICA time series (0 mean and unitary standard deviation) concatenated along the time direction. To concatenate HCP rs-fMRI data and the ones from the Traveling-subject dataset, we proceeded as follows. First, we matched the voxel coordinates of the Traveling-subject data with the group average spatial maps from the group-ICA decomposition of the HCP time series. These spatial maps were extracted from the group average analysis across all the subjects of the S1200 release and are available on the HCP website: https://www.humanconnectome.org/study/hcp-young-adult. Because the spatial maps are in a gray-ordinate CIFTI file [55], we extracted the *xyz* coordinates in a standard stereotaxic space MNI152 by using a *mid-thickness* surface file for the surface vertices and the coordinate transformation matrix included in the CIFTI file. Next, we extracted the time series from the Traveling-subject data corresponding to the same *xyz* coordinates of the aforementioned spatial map in Matlab by using the ROI Signal Extractor provided by the toolbox DPABI [59]. Finally, the HCP group average spatial map allowed us to obtain the estimated 50-dimensional ICs for the Traveling-subject data from the extracted time series. To note, once the Traveling-subject rs-fMRI time series were reduced to 50 ICs, they matched the spatial dimension of the HCP data used to infer the HMM in [33]. Finally, to train our HMM, we used the publicly available toolbox HMM-MAR (https://github.com/OHBA-analysis/HMM-MAR) [60]. We inferred *N* = 50 models from random initializations, multiple priors, and different combinations of the available datasets. Specifically, we inferred *N*_1200_ = 28 models inferred on time series from the 1200-subject HCP release only with random priors, *N*_820_ = 14 models inferred on the 820-subject HCP release (a subset of the 1200-subject release, which was used in the original work on the HMM-derived hierarchical organization of brain states [33]) with random priors, and *N_TS_* = 6 models inferred on the 9 subjects of the Traveling-Subject dataset using the best model inferred from HCP data only one as a prior, so that *N*_1200_ + *N*_820_ + *N_TS_* = 50).

### Model Selection

To select the model that best fits the data, we computed the free energy for each of the fifty different models that were inferred. The free energy provides a bound on the log-evidence for any model [61], and can be derived as the sum of the model average log-likelihood, the negative entropy and the Kullback–Leibler divergence [62]. Because the data sets have different sizes (HCP_1200_, HCP_820_, and Traveling-subject only), we corrected the free energy according to the size of the dataset used for the model inference in order to compare different models fairly (Fig. S1). We ranked the *N* = 50 models inferred in this study based on their free energy, and chose the one minimizing this quantity. Further details and matrices of notable models different from the best one can be found in Fig. S4. Since the MP Difference definition strongly relies on the notion of metastate, we also verified that the selected model presented a sufficiently marked 2-metastate structure. Therefore, we computed for each model the Euclidean distance from the ideal FO Correlation Matrix (Fig. S1). Mathematically, this distance is defined as:

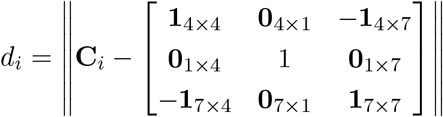

for *i* = 1, …, 50, where **C**_*i*_ if the FO Correlation Matrix of model *i*, **1** is a matrix of all ones, **0** is a zero matrix, and ‖ · ‖ denotes the Euclidean norm. The model that we have used in this study (Fig. 2) is not only the one with the lowest free energy, but also the one with and the smallest *d_i_*. Thus, our model fits the data the best and simultaneously embodies a pronounced 2-metastate structure.

Finally, to verify the robustness of our model when applied to time series other than the HCP data, we applied the HCP-trained HMM to autoregressive data (Supplementary Text 1). We found that this control analysis yields state time courses that spend most of the time on state 5 (some examples of state time courses are illustrated in Fig. S7). Moreover, state 5 is not only uncorrelated to all other states in our model, but has previously been found to be associated to motion artifacts in HCP data [33]. This result supports the robustness of our results in regards to application of our HCP-trained model to the Traveling-subject time series.

### FO Correlation Matrix and Fingerprints Computation

By applying (i.e., decoding) an HMM to a dataset with multiple subjects, we obtain the state time courses for each subject, from which it is possible to compute the vector of the Fractional Occupancy (FO) of every state for each subject. Stacking such vectors in a matrix yields the FO Matrix **R**, which is a *(no. of subjects)* × *(no. of states)* matrix that encodes state dynamics similarities across subjects. Each element **R**_*ij*_ of this matrix denotes the fraction of time spent by subject *i* in state *j*. Further, by taking the pairwise correlation of the columns of the FO Matrix **R**, we obtain the *(no. of states)* × *(no. of states)* FO Correlation Matrix

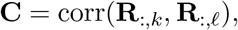

where **R**_:*,k*_ denotes the column vector of the FO of all subjects for the *k*-th state.

To compare different scanning sessions for different subjects and different factors, we compute the Metastate Profile (MP) Differences and the Fractional Occupancy Correlations across different runs. To derive these metrics, we first construct the MP matrix, whose entry (*i, k*) represents the FO of the second metastate (states 6 to 12) minus the FO of the first metastate (states 1 to 4) for the subject *i* during the scanning session *k*. Formally, given the FO Matrix **R** for the run *k*, MP_*i,k*_ is computed as follows:

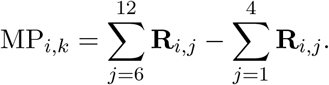

Then, the MP Difference between run *k*_1_ for subject *i*_1_ and run *k*_2_ for subject *i*_2_ reads as

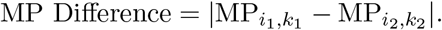

Instead, the FO Correlation between run *k*_1_ for subject *i*_1_ and run *k*_2_ for subject *i*_2_ is defined as

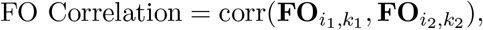

where **FO**_*i,k*_ denotes the 11-dimensional column vector of the FOs of all 12 states minus state 5 for subject *i* and run *k*.

It is worth noting that exploiting and comparing the two metrics defined above gives us a remarkable advantage with respect to utilizing only the model’s TPM. Namely, because of the stochastic nature of the model inference, we are able to avoid the non-uniqueness issue of the TPM and, at the same time, to reliably capture the temporal characteristics of the state time courses. Finally, as evident not only by their distributions of values in Fig. 4, but also from the results in Fig. 5, the two brain dynamics fingerprints employed in this study provide unique information.

### Subject Classification Using Brain Dynamics Fingerprints

To support our findings, and the robustness of the subject-specific fingerprints to data heterogeneity, we used machine learning on these fingerprints to perform subject-level classification. Specifically, individual subjects from the Traveling-subject dataset were classified based on their Metastate Profiles and Fractional Occupancies. Our simple classification experiment revealed that subject classification by means of fingerprints based on brain dynamics is possible, and that the accuracy of our classifier is largely above the baseline chance level. We detail the procedure hereafter.

For each scanning factor, we trained a logistic regression classifier – which minimizes the cross-entropy loss – with the `scikit-learn` machine learning package [63] in Python 3 with the following parameters: default L2 penalty, default L-BFGS-B algorithm [64], and ‘multi class’ option set to ‘multiomial’. The classification task is repeated multiple times by splitting the data into different training and validation datasets as follows. We repeat the training and validation of the linear regression classifier for each factor attribute (e.g., for the scanner parameter, we repeat the procedure for each scanner model) by performing a leave-one-attribute-out cross-validation: we choose as validation set all the samples (i.e. fingerprints) belonging to one factor attribute, and we used as training set all the remaining samples. We found that (1) even with different scanning protocols and heterogeneous data, the classification based on brain dynamics fingerprints performs largely above the baseline chance level, and (2) the scanning factors site and day had the lowest accuracy among all the classification results, in accordance with our claim that some factors tend to make data collections more variable than others (see Table S3 and Fig. S7).

## Supporting information

**S1 Text.** Supplementary text containing further information on the control study concerning the application of HCP-trained HMM to autoregressive time series.

**S1 Fig. Models ranking based on the free energy.** We ranked the *N* = 50 models inferred in this study based on their free energy, and chose the one minimizing this quantity. To compare all the models, each free energy calculation has been adjusted to account for different dataset sizes.

**S2 Fig. Theoretical FO Correlation Matrix.** Ideal matrix used to compare different models and their capability to capture the 2-metastate structure.

**S3 Fig. Network associated with the Transition Probability Matrix of the best model used in this study.** Weighted directed graph representation of the TPM for the best model inferred in this work after thresholding to represent only the strongest transition probabilities.

**S4 Fig. Transition Probability Matrix and Fractional Occupancy Correlation Matrix for additional HMM models.** Additional matrices associated to models worth reporting that have been inferred from different dataset partitions or priors.

**S5 Fig. Transition Probability Matrix and Fractional Occupancy Correlation Matrix for the model inferred on downsampled HCP time series.** The TPM is very skewed towards only three states: 10, 11, 12. The metastate structure in the FO Correlation Matrix is not as clear as on the original timeseries, but still emergent.

**S6 Fig. Additional Metastate Profile Differences and Fractional Occupancy Correlations.** Distributions of values for MP Differences and FO Correlations, for the following scanning factors: numbers of channels per coil, manufacturers, and scanner model.

**S7 Fig. Summary of the leave-one-attribute-out cross-validation for all scanning factors.** The three bars represent the average classification accuracy across all different scanning factors, along with the standard deviation, for the Metastate Profiles (MP), the Fractional Occupancies (FO), and the combination of the two, respectively. The red dashed line indicates the baseline chance level. These results show that the personal signature of brain dynamics fingerprints emerges even with a simple logistic regression classifier.

**S8 Fig. Examples of state time courses after HMM decoding on autoregressive time series.** To provide a baseline for our study, we applied the HMM model used in this work to autoregressive data with dimension 50 × *T*, where *T* denotes the number of time points. The HMM decoding yields state time courses that stay most of the time in state 5, which is the state that is highly uncorrelated from the other 11 states and the one with the highest variance. This fact supports the goodness of fit of the inferred model, as randomized time series do not provide meaningful state time courses.

**S1 Table. Imaging protocols for rs-fMRI in the Traveling-subject dataset.** All scanning protocols and parameters of the Traveling-subject dataset for all the sites at which fMRI scans where acquired.

**S2 Table. Kolmogorov-Smirnov test statistics for MP Differences and FO Correlations.** SS: Same subject Same factor attribute. SD: Same subject Different factor attributes. DS: Different subjects Same factor attribute.

**S3 Table. Logistic regression accuracy results.** Accuracy results of a simple leave-one-attribute-out cross-validation for all scanning factors obtained through logistic regression.

## Acknowledgments

We thank the Washington University–University of Minnesota Consortium of the Human Connectome Project (WU-Minn HCP) for generating and making publicly available the HCP data. Data used in the preparation of this work were obtained from the DecNef Project Brain Data Repository (https://bicr-resource.atr.jp/srpbsts/) gathered by a consortium as part of the Japanese Strategic Research Program for the Promotion of Brain Science (SRPBS) supported by the Japanese Advanced Research and Development Programs for Medical Innovation (AMED). T. M. is supported by the National Science Foundation (NSF) (USA, grant NCS-FO-1631112), by the Army Research Office (ARO) (USA, grant ARO-71603NSYIP), by JST ERATO (Japan, grant JPMJER1801), and by AMED (Japan, grant JP18dm0307008). G.L. is supported by AMED (Japan, grant JP18dm0307002). A. C. is supported by JST ERATO (Japan, grant JPMJER1801) and by AMED (Japan, grants JP18dm0307002 and JP18dm0307008).

## Notes

https://bicr.atr.jp/decnefpro/data/

### Competing Interest Statement

The authors have declared no competing interest.

