## Supplementary Information for "Brain network dynamics fingerprints are resilient to data heterogeneity"

Tommaso Menara
Department of Decoded Neurofeedback,
ATR Computational Neuroscience Laboratories,
2-2-2 Hikaridai, Seika-cho, Soraku-gun, Kyoto, 619-0288, Japan
Department of Mechanical Engineering,
University of California Riverside,
900 University Ave, Riverside, CA, 92521, USA

Giuseppe Lisi
Nagoya Institute of Technology,
Gokiso-cho, Showa-ku, Nagoya, Aichi, 466-8555, Japan
ATR Brain Information Communication Research Laboratory Group,
2-2-2 Hikaridai, Seika-cho, Soraku-gun, Kyoto, 619-0288, Japan

Fabio Pasqualetti
Department of Mechanical Engineering,
University of California Riverside,
900 University Ave, Riverside, CA, 92521, USA

Aurelio Cortese
Department of Decoded Neurofeedback,
ATR Computational Neuroscience Laboratories,
2-2-2 Hikaridai, Seika-cho, Soraku-gun, Kyoto, 619-0288, Japan


### Supplementary Note

#### Control Analysis with Autoregressive Data

To verify the robustness of our analysis in regards to the application of the HMM model to the Traveling-subject dataset, we applied the HCP-trained HMM to autoregressive data. To assess a model AR($n$) of order $n$ fairly and prevent it from trying to disproportionately fit to the first $n$ datapoints, we discarded the first $3n$ datapoints. For this analysis, we have chosen autoregressive model of order 5. With this AR(5) model, we have generated 50 time series as dummy ICA timeseries to be used the model inference. Analogously to the real ICA timeseries, the AR time series have been normalized (zero mean and unitary standard deviation).

The decoding of our model on autoregressive time series yielded state time courses that in which state 5 is predominant in all different scanning sessions. See Fig. S8 for a few examples of the state time courses obtained from HMM decoding of the randomly permute Traveling-subject time series. It is worth noting that state 5 is the state that is mostly uncorrelated from the remaining 11 states and it is the state with the largest variance. In [1], it is demonstrated that state 5 is associated with motion artifacts in the scanner. The outcome of the HMM decoding on autoregressive data is in accordance with these observations, therefore supporting the robustness of our findings.

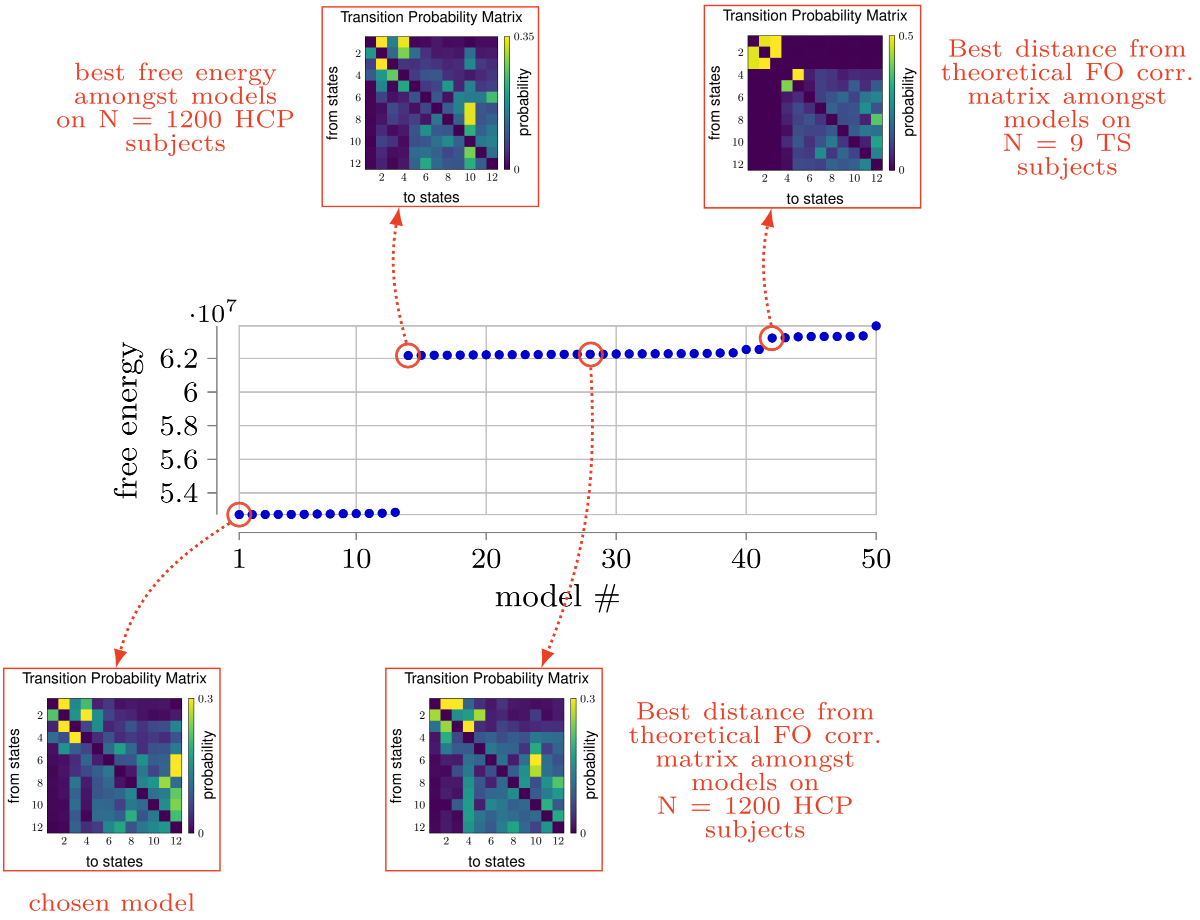


Figure S1: **Models ranking based on the free energy.**
We ranked the $N = 50$ models inferred in this study based on their free energy, and chose the one minimizing this quantity. To compare all the models ($N_{1200} = 28$ models inferred on 1200-subject HCP release, $N_{820} = 14$ models inferred on the 820-subject HCP release, and $N_{TS} = 6$ models inferred on the 9 subjects of the Traveling-Subject dataset, so that $N_{1200} + N_{820} + N_{TS} = 50$), each free energy calculation has been adjusted to account for different dataset sizes. The FO Correlation Matrix can be found in Figure S2, and further details on different models from the best one can be found in Figure S4.

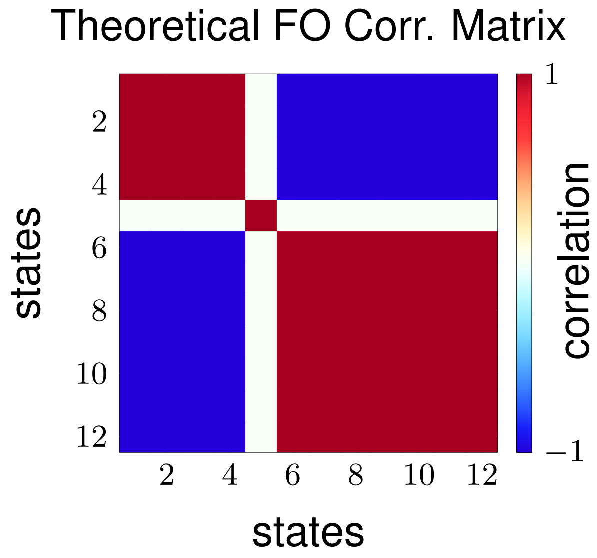


Figure S2: **Ideal FO Correlation Matrix.**
For each model inferred from our datasets, we compute the Euclidean distance between the model's FO Correlation Matrix and the ideal one. We rank our models based on this distance with the aim of selecting the model that has the most clear emergence of the 2-metastate structure.

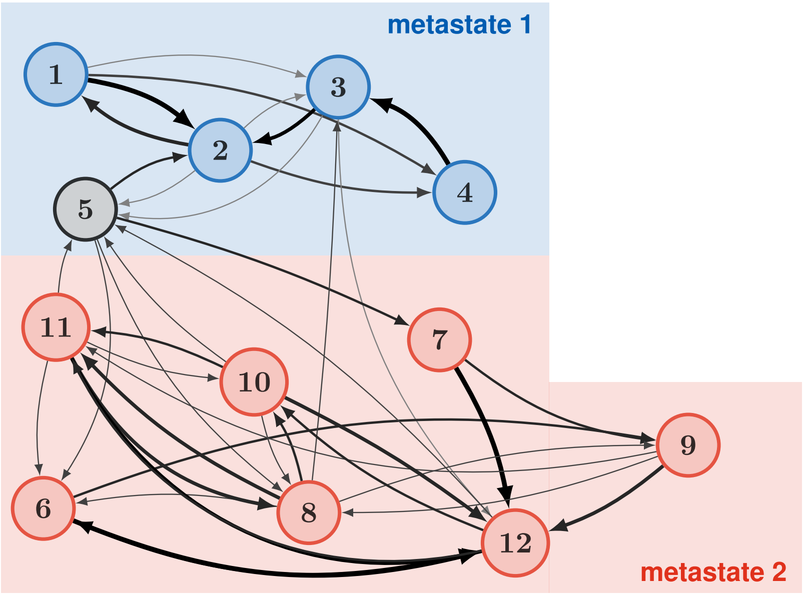


Figure S3: **Network associated with the Transition Probability Matrix of the HMM used in this study (Fig. S3A).**
The first 4 states comprise metastate 1, whereas states 6 to 11 comprise metastate 2. State 5 is mostly uncorrelated to the other states, is associated with head motion, and has the highest variance [1]. The interconnections depicted in this graph represent probabilities higher than 10% (i.e. $> 0.1$) in the HMM's Transition Probability Matrix, and their color and thickness are proportional to their magnitude.

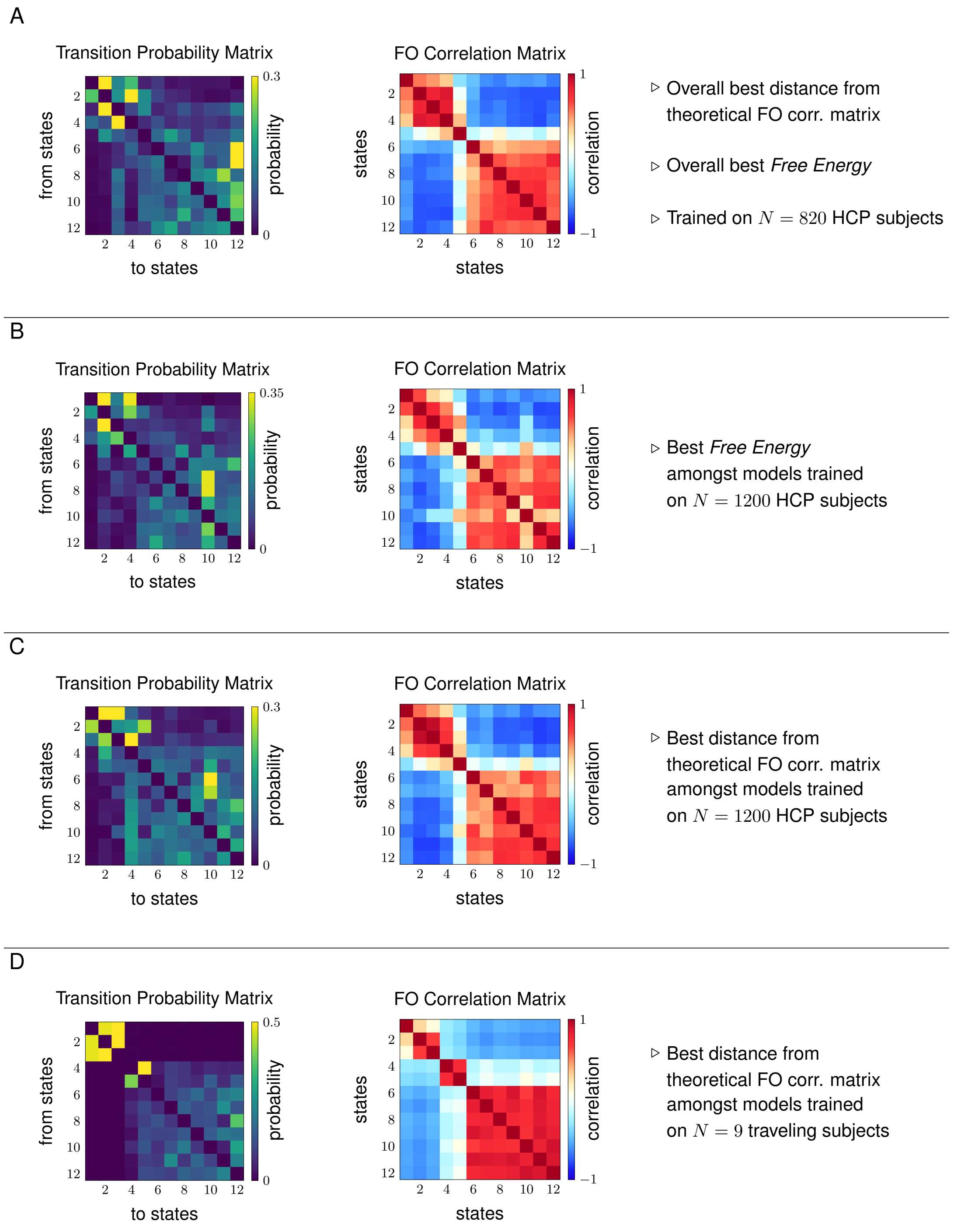


Figure S4: **Transition Probability Matrix and Fractional Occupancy Correlation Matrix for different HMM models.**
**A**, The best model among all the models trained in terms of free energy and Euclidean distance from the ideal FO Correlation Matrix. **B**, The best model in terms of free energy among all models trained on the subjects of the HCP 1200-subject distribution. **C**, The best model in terms of Euclidean distance from the ideal FO Correlation Matrix among all models trained on the subjects of the HCP 1200-subject distribution. **D**, The best model in terms of Euclidean distance from the ideal FO Correlation Matrix among all models trained on the Traveling-subject dataset by using the model **A** as a prior. Notice that, while the model in this last panel displays very distinct metastate separation in the TPM matrix, such a matrix is not irreducible (i.e. there does not exist a path connecting the two groups of states), making it not suitable to represent any biological system.

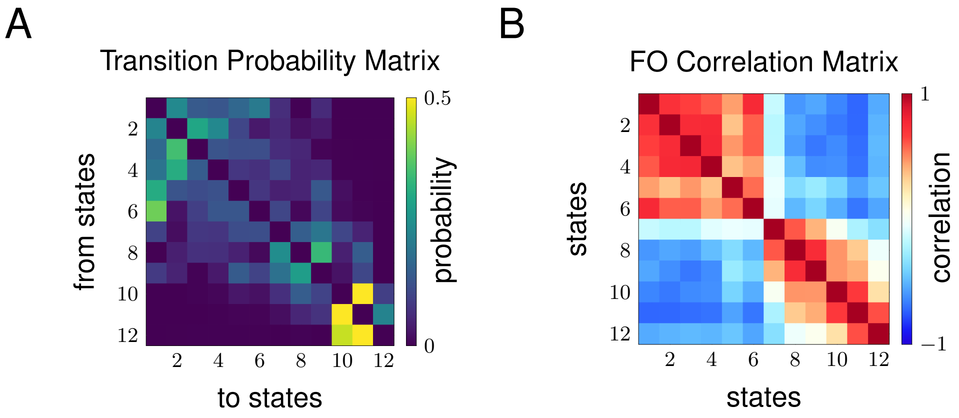


Figure S5: **Best Model inferred on downsampled HCP timeseries.**
(A) Transition probability matrix for the model inferred on downsampled HCP time series. The TPM is very skewed towards only three states: 10,11, 12. (B) FO Correlation Matrix for the model inferred on downsampled HCP time series. The metastate structure is not as clear as on the original timeseries, but still emergent.

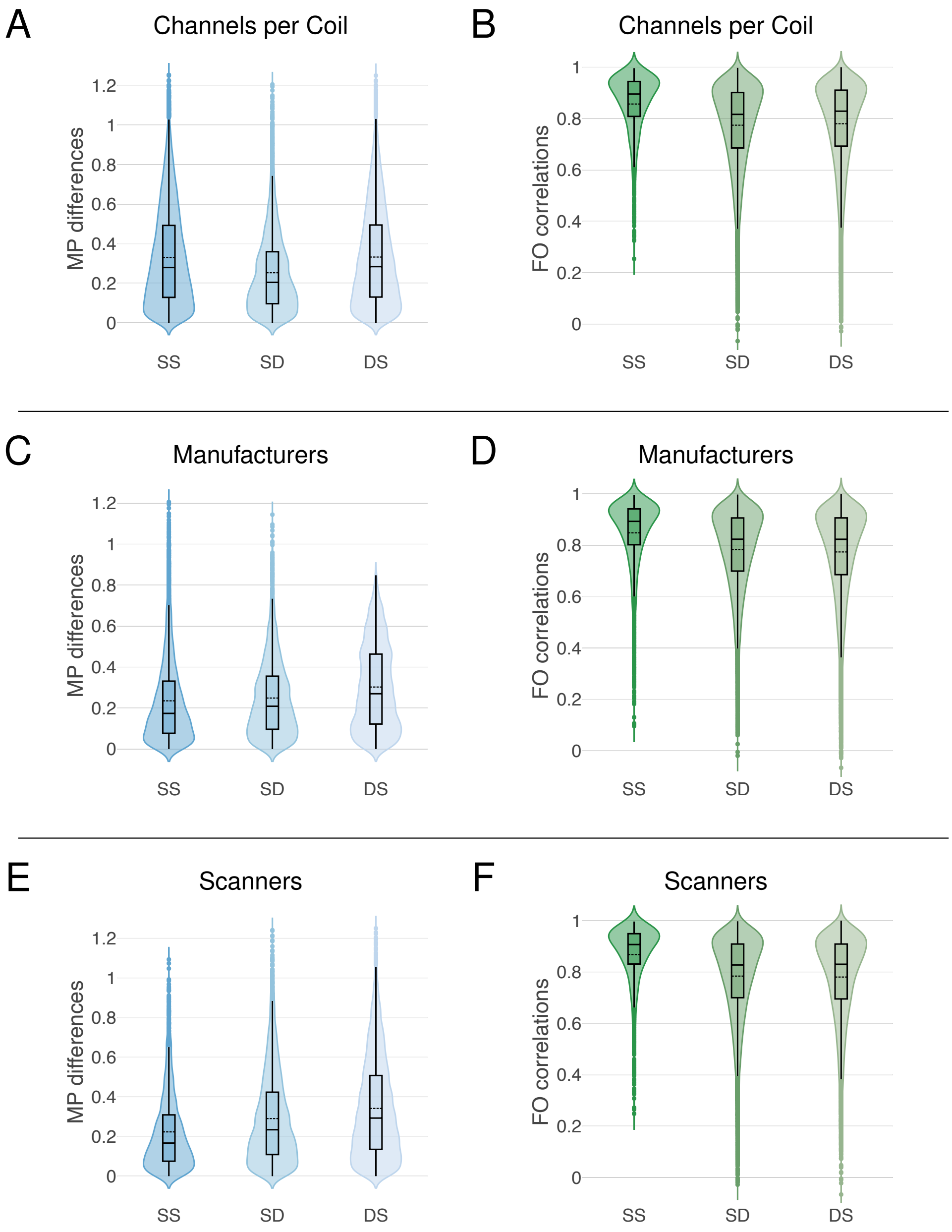


Figure S6: **distributions of values for MP Differences and FO Correlations, for the factors: numbers of channels per coil, manufacturers, and scanner model.**
In panels **A** to **F**, the set SS comprises the MP Differences (resp., FO Correlations) computed for each subject within the same factor attribute, and the SS distribution displays these values for all subjects; the set SD consists of the MP Differences (resp., FO Correlations) computed for each subject across different attributes of the same factor, and the SD distribution displays these values for all subjects; finally, the set DS consists of the MP Differences (resp., FO Correlations) computed across all subjects within the same factor attribute, and the DS distribution displays these values for all attributes of the same factor. Further, for all the distributions, the black dashed lines illustrate the mean. The difference between SS and DS distributions in panel **A**, and the difference between SD and DS distributions in panel **F**, are not statistically significant (see also Table 1 in the main text).

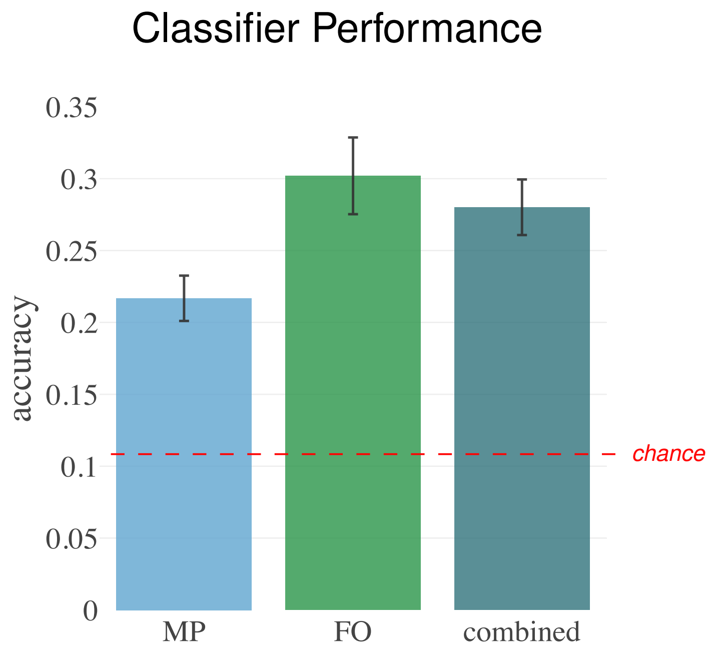


Figure S7: **Summary of the leave-one-attribute-out cross-validation for all scanning factors.**
The three bars represent the average classification accuracy across all different scanning factors, along with the standard deviation, for the Metastate Profiles (MP), the Fractional Occupancies (FO), and the combination of the two, respectively. The red dashed line indicates the baseline chance level. These results show that the personal signature of brain dynamics fingerprints emerges even with a simple logistic regression classifier.

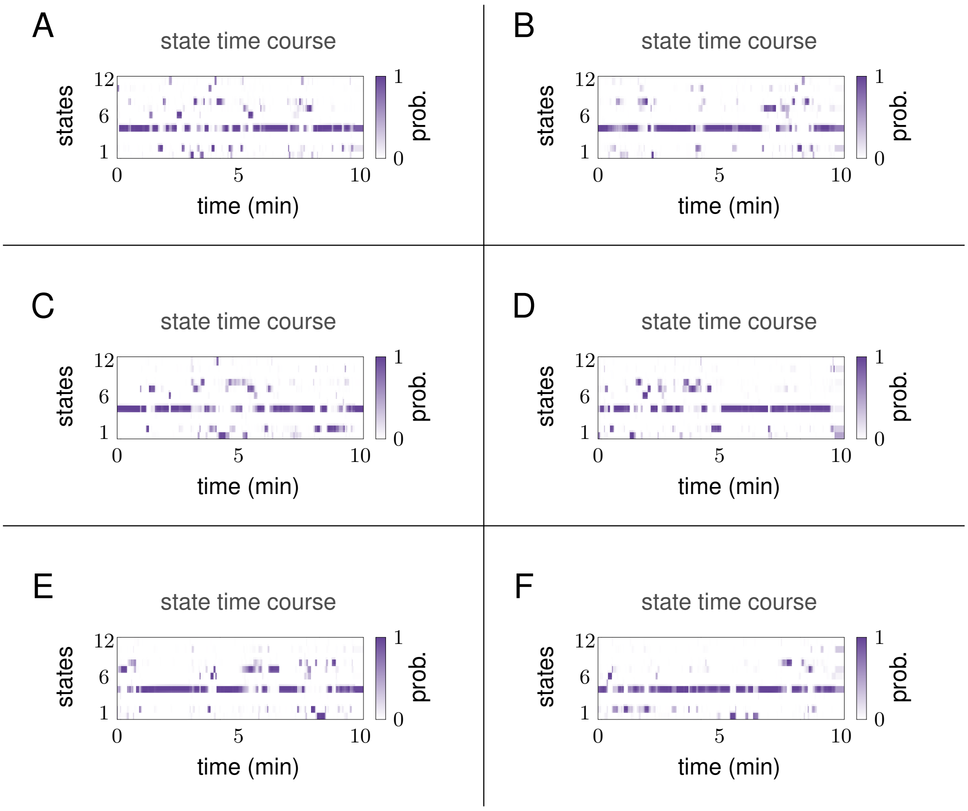


Figure S8: **Examples of state time courses after HMM decoding on AR(5) time-series.**

**A-F**, To provide a baseline for our study, we applied the HMM model used in this work to autoregressive data with dimension $50 \times T$, where $T$ denotes the number of time points. The HMM decoding yields state time courses that stay most of the time in state 5, which is the state that is highly uncorrelated from the other 11 states and the one with the highest variance. This fact supports the goodness of fit of the inferred model, as randomized time series do not provide meaningful state time courses.

Table S1: Imaging protocols for resting-state fMRI in the Traveling-subject dataset.

| Site | ATT | ATV | COI | HUH | HKH | KPM | SWA | KUT | KUS | UTO | YC1 | YC2 |
|---|---|---|---|---|---|---|---|---|---|---|---|---|
| 1. Scanner Manufacturer | Siemens | Siemens | Siemens | GE | Siemens | Philips | Siemens | Siemens | Siemens | GE | Philips | Philips |
| 2. Scanner Model | TimTrio | Verio | Verio | Signa HDxt | Spectra | Achieva | Verio | TimTrio | Skyra | MR750W | Achieva | Achieva |
| 3. Magnetic Field Strength | 3.0 T | 3.0 T | 3.0 T | 3.0 T | 3.0 T | 3.0 T | 3.0 T | 3.0 T | 3.0 T | 3.0 T | 3.0 T | 3.0 T |
| 4. No. Channels per Coil | 12 | 12 | 12 | 8 | 12 | 8 | 12 | 32 | 32 | 24 | 8 | 8 |
| 5. Field-of-view (mm) | $212 \times 212$ | $212 \times 212$ | $212 \times 212$ | $212 \times 212$ | $212 \times 212$ | $212 \times 212$ | $212 \times 212$ | $212 \times 212$ | $212 \times 212$ | $212 \times 212$ | $212 \times 212$ | $212 \times 212$ |
| 6. Matrix | $64 \times 64$ | $64 \times 64$ | $64 \times 64$ | $64 \times 64$ | $64 \times 64$ | $64 \times 64$ | $64 \times 64$ | $64 \times 64$ | $64 \times 64$ | $64 \times 64$ | $64 \times 64$ | $64 \times 64$ |
| 7. No. of Slices | 40 | 39 | 40 | 35 | 35 | 40 | 40 | 40 | 40 | 40 | 40 | 40 |
| 8. No. of Volumes | 240 | 240 | 240 | 240 | 240 | 240 | 240 | 240 | 240 | 240 | 240 | 240 |
| 9. In-plane Resolution (mm) | $3.3125 \times 3.3125$ | $3.3125 \times 3.3125$ | $3.3125 \times 3.3125$ | $3.3125 \times 3.3125$ | $3.3125 \times 3.3125$ | $3.3125 \times 3.3125$ | $3.3125 \times 3.3125$ | $3.3125 \times 3.3125$ | $3.3125 \times 3.3125$ | $3.3125 \times 3.3125$ | $3.3125 \times 3.3125$ | $3.3125 \times 3.3125$ |
| 10. Slice Thickness (mm) | 3.2 | 3.2 | 3.2 | 3.2 | 3.2 | 3.2 | 3.2 | 3.2 | 3.2 | 3.2 | 3.2 | 3.2 |
| 11. Slice Gap (mm) | 0.8 | 0.8 | 0.8 | 0.8 | 0.8 | 0.8 | 0.8 | 0.8 | 0.8 | 0.8 | 0.8 | 0.8 |
| 12. TR (ms) | 2.5 | 2.5 | 2.5 | 2.5 | 2.5 | 2.5 | 2.5 | 2.5 | 2.5 | 2.5 | 2.5 | 2.5 |
| 13. TE (mm) | 30 | 30 | 30 | 30 | 30 | 30 | 30 | 30 | 30 | 30 | 30 | 30 |
| 14. Total Scan Time (min:s) | 10 : 00 | 10 : 00 | 10 : 00 | 10 : 00 | 10 : 00 | 10 : 00 | 10 : 00 | 10 : 00 | 10 : 00 | 10 : 00 | 10 : 00 | 10 : 00 |
| 15. Flip Angle (deg) | 80 | 80 | 80 | 80 | 80 | 80 | 80 | 80 | 80 | 80 | 80 | 80 |
| 16. Slice Acquisition Order | Ascending | Ascending | Ascending | Ascending | Ascending | Ascending | Ascending | Ascending | Ascending | Ascending | Ascending | Ascending |
| 17. Phase Encoding | PA | PA | AP | PA | PA | AP | PA | PA | AP | PA | AP | AP |
| 18. Eyes Closed/Fixate | Fixate | Fixate | Fixate | Fixate | Fixate | Fixate | Fixate | Fixate | Fixate | Fixate | Fixate | Fixate |
| 19. Field Map | yes | yes | yes | - | - | yes | yes | yes | yes | yes | yes | - |

Table S2: Kolmogorov-Smirnov test statistics for MP Differences and FO Correlations

| | MP Differences | | | FO Correlations | | |
|---|---|---|---|---|---|---|
| Parameter | SS vs SD | SS vs DS | SD vs DS | SS vs SD | SS vs DS | SD vs DS |
| 1. Site | 0.2808 | 0.1756 | 0.1552 | 0.5535 | 0.4549 | 0.1416 |
| 2. Day | 0.4438 | 0.2183 | 0.2468 | 0.4708 | 0.2424 | 0.2745 |
| 3. Phase | 0.0428 | 0.1904 | 0.154 | 0.044 | 0.2074 | 0.1687 |
| 4. Channels/Coil | 0.1422 | 0.0078 | 0.1472 | 0.2513 | 0.2168 | 0.0361 |
| 5. Manufacturer | 0.0735 | 0.1836 | 0.1419 | 0.2184 | 0.2156 | 0.0257 |
| 6. Scanner | 0.1454 | 0.231 | 0.0929 | 0.2616 | 0.2686 | 0.0117 |

Table S3: Logistic regression accuracy results

| Parameter | MP | FO | combined |
|---|---|---|---|
| 1. Site | 0.2096 | 0.2944 | 0.29259 |
| 2. Day | 0.1889 | 0.2657 | 0.2509 |
| 3. Phase | 0.2222 | 0.2871 | 0.2723 |
| 4. Channels/Coil | 0.2341 | 0.3008 | 0.2837 |
| 5. Manufacturer | 0.2217 | 0.3416 | 0.3074 |
| 6. Scanner | 0.2243 | 0.3220 | 0.2736 |
| Mean | 0.2168 | 0.3020 | 0.2801 |
| Standard Deviation | 0.0158 | 0.02668 | 0.0193 |
